# Integration of sensory and cortical information in the brainstem during mastication in mice

**DOI:** 10.1101/2025.07.29.667274

**Authors:** Sophia Dubois, Dominic Falardeau, Ohini Yanis Sanvi, Dorly Verdier, Arlette Kolta

## Abstract

Mastication is a vital function that relies on precise synchronization among multiple brainstem regions, known as being part of a central pattern generator (CPG). Movements can be triggered either by stimulating a sensory-motor region called the cortical masticatory area (CMA), well-documented in various species but not yet formally identified in mice, or by stimulating the oro-facial sensory primary afferents which fibers form the trigeminal tract (Vtr). However, it’s unclear whether these different inputs activate distinct components of the CPG or converge on the same. This study aims at mapping brainstem areas activated by cortical and sensory inputs using immunohistochemistry against the cellular activity marker, c-Fos and Ca^2+^-imaging, respectively. Optogenetic stimulation of the cortical masticatory area (CMA) in awake, head-fixed mice reliably induced rhythmic jaw movements (RJMs) and increased c-Fos expression in multiple brainstem regions, with strongest activation in the peritrigeminal area (PeriV) and parvocellular reticular formation (PCRt) ventral to the trigeminal motor nucleus (NVmt). In contrast, in vitro electrical stimulation of trigeminal sensory afferents (Vtr) predominantly activated neurons and astrocytes in the main sensory nucleus (NVsnpr), the dorsal area of PeriV, adjacent to it, and PCRt. The areas containing the highest numbers of activated cells differed clearly according to the type of inputs and overlapped only in the PCRt, ventral and slightly medial to the trigeminal motor nucleus and the most dorso-medial part of PeriV. These findings demonstrate that cortical and sensory inputs take part in distinct components of the brainstem masticatory circuitry, with PCRt emerging as a point of convergence and provide new insights into the components of the CPG of mastication.

## Introduction

Mastication, a vital function that involves preparing food for digestion, is undoubtedly a key component of feeding behavior. Far from being a stereotyped sequence of jaw-closing and -opening movements, mastication relies on sensory and cortical inputs to adapt the movements to the properties of the food bolus. Using decerebrate and paralyzed animals, Dellow and Lund (1971) were the first to demonstrate that like other rhythmic movements such as breathing and locomotion, mastication relies on a specific neuronal network known as a central pattern generator (CPG) located in the brainstem in the case of mastication (Feldman and Grillner, 1983; Lund, 1991; Falardeau et al., 2023). Further work from the group of Matsuya, (Kogo et al., 1996; Tanaka et al., 1999) using an isolated brainstem preparation suggested that the masticatory CPG was confined to the pontine area of the brainstem defined rostro-caudally by the rostral limits of the trigeminal (NVmt) and facial (NVII) motor nuclei and medio-laterally by the column of the last-order interneurons surrounding the NVmt and the trigeminal sensory complex formed at this level by the trigeminal main sensory nucleus (NVsnpr) and the rostral part of the spinal trigeminal nucleus (NVspo). The areas dorsal, lateral and medial to NVmt named the supra-(SupV), inter-(IntV) and juxta-trigeminal (JuxtV) areas, respectively, form collectively the peritrigeminal area (PeriV). While the area ventral to NVmt, which also extends caudally, is the parvocellular reticular formation (PCRt). Besides a small proportion of SupV neurons, intrinsic bursting properties were only found in neurons of the dorso-medial part of NVsnpr (Sandler et al., 1998; Bourque and Kolta, 2001; Brocard et al., 2006), which led to the proposal that it forms the rhythmogenic core of the masticatory CPG (Tsuboi et al., 2003; Brocard et al., 2006, Morquette et al., 2015).

In *in vivo* animal models, rhythmic jaw movements (RJMs) resembling mastication can be elicited by sustained repetitive stimulation of a cortical area named the cortical masticatory area (CMA) or of the trigeminal sensory afferents (Dellow and Lund, 1971), which both project to components of the brainstem area thought to contain the masticatory CPG. The aim of this study was to examine whether these inputs engage different components or converge on the same areas to better understand the circuitry underlying mastication. The CMA has been extensively studied in many species but is still not yet clearly defined in mice; although Lindsay *et al*. pointed at its existence in a study mapping corticobulbar projections (Mercer Lindsay et al., 2019). In rabbits, c-Fos expression in trigeminal areas has been used to map neurons activated by stimulation of the CMA (Athanassiadis et al., 2005). Here, a similar approach was used to describe the distribution of neurons activated by stimulation of the CMA in mice but could not be used to map distribution of cells activated by sensory inputs, since it is difficult to activate these specifically and strongly in-vivo. Thus, we used Ca^2+^-imaging in *in vitro* slice preparations to map the distribution of cells responding to electrical stimulation of the portion of the Vth tract containing the fibers of sensory afferents most needed for mastication (periodontal and muscle spindle afferents). Our results confirmed the existence of a circumscribed CMA in the mice and revealed distinct patterns of cellular activation. CMA stimulation broadly increased neuronal activity across multiple brainstem regions, including the NVsnpr with pronounced activation primarily in JuxtV and PCRt. In contrast, sensory stimulation predominantly revealed cellular activation in NVsnpr, IntV, SupV, and PCRt. These observations suggest that PCRt may serve as a critical convergence hub for both cortical and sensory inputs.

## Methods

All procedures followed the guidelines of the Canadian Institutes of Health Research and were approved by the University of Montreal Animal Care and Use Committee. A comprehensive description of the materials and methods can be found in the Supplemental Materials and Methods.

## Results

### Optogenetic stimulation of the CMA in vivo in awake head-fixed animals induces RJMs

Using whole cell patch clamp recordings of layer 5 pyramidal neurons, we first examined in *in vitro* cortical slices from VGlut2-ChR2 mice, the effects of photostimulation to ensure that it efficiently activates pyramidal neurons. Indeed, as depicted in **Figure 1A**, blue light (440 nm; top trace), but not red light (561 nm; bottom trace) induced sustained firing in the recorded neurons (n=5; N=2). Then, to elicit RJMs with optogenetic stimulation of corticobulbar fibers, we implanted optic fibers in VGlut2-ChR2 mice (**Figure 1B**). As in rats, we found an anterior area (coordinates: 2.5mm AP, 2.0mm L and 0.75mm D relative to Bregma) in the motor cortex and a posterior area (coordinates: 0mm AP, 3.5mm L and 2.0mm D relative to Bregma) in the granular and agranular insular posterior cortices (**Figure 1C**). We confirmed ChR2 expression and optic fiber implantation location by performing immunohistochemistry on the brain sections of the tested animals to reveal the GFP-stained cells (arrows) and their NeuN-stained nuclei (arrowheads) as well as the lesion (*) made by the optic fiber (**Figure 1D**). Optogenetic stimulation at 40Hz (for 3 sec) induced stereotypical RJMs, as shown in **Figure 1E** and **1F** for anterior and posterior areas, respectively. For a-area stimulation, the movements began with the stimulation and continued throughout its duration until the end. Conversely, for p-area, movements sometimes persisted for 1-2 seconds after the end of stimulation. We then assessed whether RJMs maintained a relatively stable frequency over the course of the 30-minutes stimulation period. The frequency of RJMs in mice stimulated in both the a-area and p-area combined remained consistent from start to end (Start=11.17±0.74Hz vs End=10.95±0.45Hz, *P*=0.6875>0.05, two-tailed Wilcoxon matched-pairs signed rank test, N=6; **Figure 1G**). Overall, we established a robust method to induce RJMs for the first time in awake and head-fixed mice using optogenetics, demonstrating consistent results over time.

**Figure 1:**
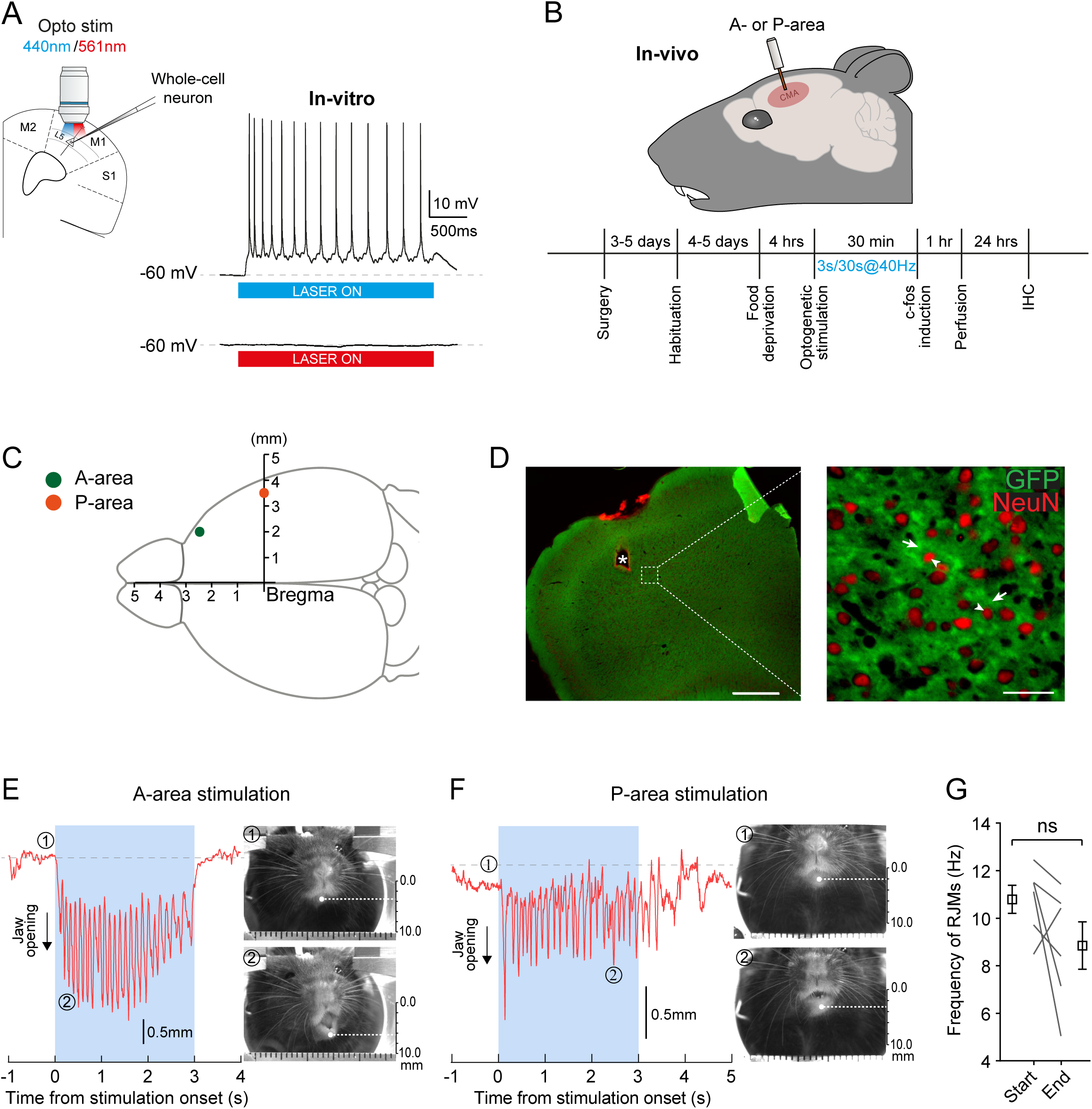
*In vivo* optogenetic stimulation of the CMA induces RJMs. (A) *In vitro* validation of the optogenetic stimulation using patch-clamp recording of a layer V cortical neuron that responds to blue light (440 nm laser, top trace) but not to red light (561 nm laser, bottom trace). **(B) Left** Timeline of the *in vivo* experiments. **(C)** Dorsal view showing coordinates of the anterior (green) and posterior (orange) cortical areas stimulated. **(D)** Immunohistochemistry against GFP (green) and NeuN (red) in VGlut2-ChR2 mice to confirm the presence of ChR2 in the stimulated area. The asterisk indicates the placement of the optic fiber. The scale bars in the low magnification and high magnification images are 500 and 200 µm, respectively. Examples of the vertical component of RJMs kinematics extracted from DeepLabCut traces induced by stimulation (40 Hz, 2.5 ms pulse, 3-second duration) in the A-area **(E)** and the P-area **(F)** of the CMA in VGlut2-ChR2 mice. Pictures on the right show the vertical position of the lower jaw at its closed state (1) prior to the optogenetic stimulation of the CMA and at its opened state (2) during the optogenetic stimulation of the CMA. The scale bars on the right are 10 mm. **(G)** The frequency of the RJMs showed no significant change between the start and end of the experiment (Start=11.17±0.74Hz vs End=10.95±0.45Hz, *P*=0.6875>0.05, two-tailed Wilcoxon matched-pairs signed rank test, N=6).

### Optogenetic stimulation of the CMA increases c-Fos expression in many brainstem areas, with the strongest effect in PeriV

Brainstem c-Fos-stained cells were mapped, as described in the methods, for 6 mice that were subjected to optogenetic stimulation of the CMA (Stimulated) and for 5 mice that were implanted but unstimulated (Control). The paired interleaved scatter plot in **Figure 2A** displays data from both control (empty circles) and stimulated (blue circles) animals, illustrating the quantitative distribution of c-Fos staining among the brainstem nuclei and regions. The *P*-values for all regions are presented in **Table 1**. Since projections from the CMA are predominantly contralateral, we investigated whether the position of the fibers influenced activation patterns. To this end, we compared the ipsilateral (purple) and contralateral (orange) sides to identify any significant differences. As shown in **Figure S1A**, no significant changes were observed in any of the regions analyzed. We then examined whether stimulation of the a-area led to changes in c-Fos-stained cell counts compared to stimulation of the p-area (**Figure S1B,** gray and mauve circles, respectively). Although the number of mice implanted in the p-area was still limited, no differences were detected between the two groups across all regions. We then performed a density mapping of all brainstem sections to pinpoint the hotspot of c-Fos-stained cells. For analysis purposes, data for regions expanding on multiple sections (as shown in **Figure 2B and 2C**) were combined in the plots (**Figure 2A and S1A, B)** and in **Table 1**. Overall, several regions showed an increase in c-Fos-stained cells in stimulated animals relatively to control, including the PCRt, JuxtV, SupV, IntV, NVmt, the trigeminal accessory motor nucleus (NVac), and the NVsnpr.

**Figure 2:**
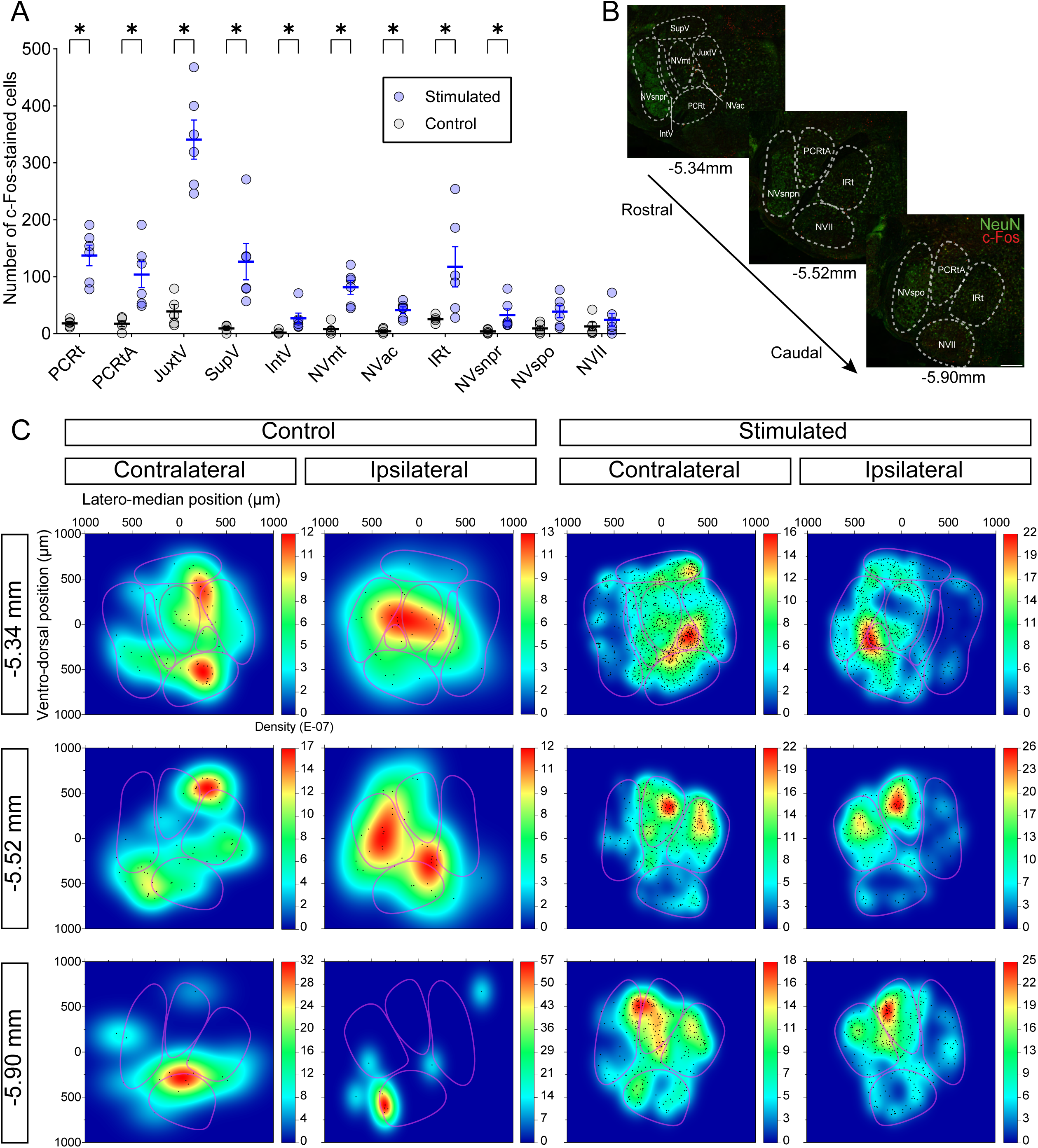
Distribution of c-Fos-stained cells in the brainstem induced by optogenetic stimulation of the CMA. **(A)** The paired interleaved scatter plots give a summary of c-Fos-stained cells distribution across the brainstem regions. Cortical stimulation was consistently delivered on the right side of the brain. Descriptive analysis and *P*-values are found in **Table 1**. **(B)** Three main rostro-caudal levels in the brainstem were assessed for density mapping: rostral (-5.34 mm), middle (-5.52 mm) and caudal (-5.90 mm) relative to bregma. The scale bar is 500 µm. **(C)** Density mapping revealed clusters of c-Fos-stained cells at specific hotspots across the 3 assessed levels. Each map represents a superimposition of data from 5 control and 6 stimulated animals. Note that, to facilitate the detection of hotspots in both control and stimulated groups across different regions, different scale bars are used.

**Table 1.** Descriptive statistics of quantification of c-Fos-stained cells per region.

|  | Control<br>(number of c-Fos-stained cells)<br>(N=5) | Stimulated<br>(number of c-Fos-stained cells)<br>(N=6) | Adjusted<br>*P*-value |
| --- | --- | --- | --- |
| PCRt | 18 ± 2 | 149 ± 12 | **0.047** |
| PCRtA | 17 ± 4 | 86 ± 15 | **0.047** |
| JuxtV | 39 ± 9 | 329 ± 29 | **0.047** |
| SupV | 9 ± 2 | 136 ± 24 | **0.047** |
| IntV | 2 ± 1 | 30 ± 4 | **0.047** |
| NVmt | 8 ± 3 | 74 ± 10 | **0.047** |
| NVac | 4 ± 2 | 45 ± 3 | **0.047** |
| IRt | 25 ± 2 | 90 ± 19 | **0.047** |
| NVsnpr | 4 ± 1 | 36 ± 10 | **0.047** |
| NVspo | 9 ± 3 | 35 ± 9 | 0.101 |
| NVII | 13 ± 5 | 15 ± 5 | 0.537 |

Values are mean ± SEM. N = number of mice Adjusted *P*-value was calculated using Mann-Whitney U tests due to non-normality distributed data for c-Fos data with Holm-Šídák correction for multiple comparisons.

At the level of the NVmt (-5.34 mm relative to Bregma; **Figure 2C, top**), density maps on the contralateral side revealed that c-Fos-positive cells were primarily distributed in the PCRt, NVac, NVmt, and JuxtV. A similar pattern was observed ipsilaterally. Control experiments revealed clusters of c-Fos-positive cells localized partially in the PCRt, JuxtV, and NVmt on the contralateral side, and partially around the JuxtV, NVac, and NVmt on the ipsilateral side.

Similar results were obtained at the level of the medial pons (-5.52 mm relative to Bregma; **Figure 2C, middle**) and close to the medulla (-5.90 mm relative to Bregma; **Figure 2C, bottom**). In both levels, c-Fos-stained cells were predominantly located in the dorsomedial part of the brainstem, with density hotspots centered around the PCRtA and IRt. Density maps from control experiments at the medial pons showed sparse staining at the junction of the IRt and PCRtA on the contralateral side, while on the ipsilateral side, staining was primarily localized within the IRt. In the medulla, control results differed from the experimental condition, with staining hotspots observed in the PCRtA on the contralateral side at both examined locations.

Thus, optogenetic stimulation of the CMA led to a marked increase in c-Fos expression in several brainstem regions. These increases were more important for rostral areas analyzed, slightly higher on the contralateral side and highest in JuxtV. These results highlight a lateralized and region-specific activation pattern within the masticatory CPG circuitry.

### In vitro stimulation of the sensory inputs generates repetitive calcium transients in neurons and astrocytes of the NVsnpr and PeriV

Based on the above results, we pursued experimentation using calcium imaging techniques in *in vitro* brainstem slices at the level of the NVmt, where the highest increases in c-Fos were seen, to map areas responding most strongly to stimulation of sensory inputs conveyed by primary afferent axons forming the trigeminal tract (Vtr). This approach makes it possible to visualize both neuronal and astrocytic calcium responses to stimulation.

Expression of the calcium indicator GCaMP6f by brainstem neurons was confirmed in two lines of transgenic mice expressing it under the control of VGlut2 (**Figure 3A**) and Thy1 promoters (**Figure 3B**). To detect astrocytic responses simultaneously with neuronal responses, we added Fluo8L-AM, which is a calcium indicator primarily taken up by astrocytes when added to the bath and SR-101 was also used to label astrocytes specifically (**Figure 3C, middle**). Neurons and astrocytes in the dorsal NVsnpr (**Figure 3E**, traces 1-2) and PeriV (**Figure 3E**, traces 3-4) responded to glutamate (500 µm, 30s) applications, confirming the functionality of our two calcium indicators (GCaMP6f and Fluo8L).

**Figure 3:**
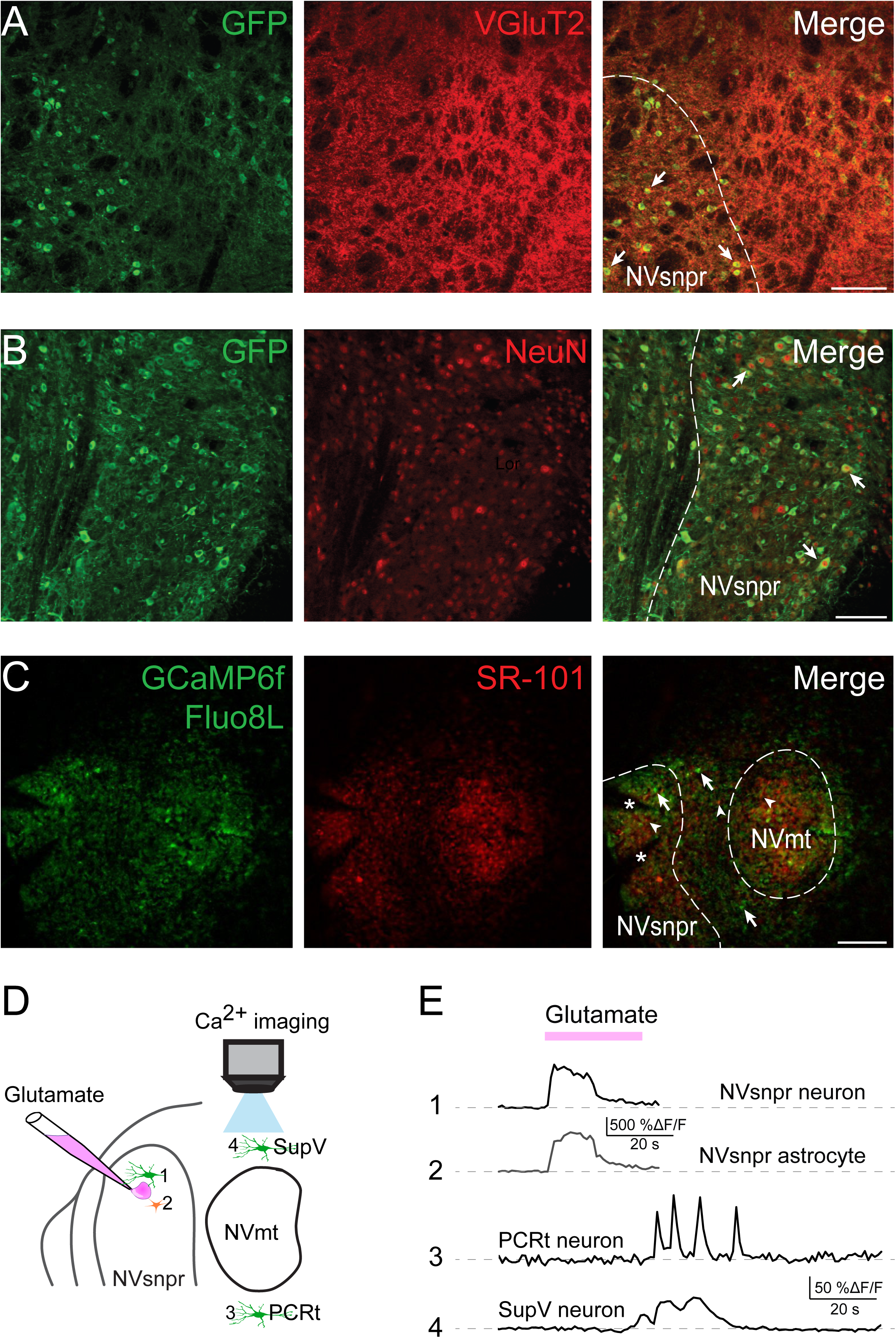
Validation of the *in vitro* model to assess brainstem Ca^2+-^imaging. The calcium indicator GCaMP6f is expressed under the control of the neuronal promoters Thy1 and VGlut2 in two separate strains of mice. **(A)** The green fluorescent protein (GFP, green) used to identify GCaMP6f is expressed in NVsnpr neurons cytoplasm (left). The overlay of GFP and VGlut2 labeling shows a few neurons expressing both proteins (right, arrows). **(B)** GFP (green) in NVsnpr neurons cytoplasm allows identification of GCaMP6f, in a Thy1-GCaMP6f mouse. NeuN (red) identifies nuclei of neurons in the slice. Co-labeling of NeuN and GFP allowed identification of projection neurons expressing GCaMP6f (arrows). **(C) Left**: Photomicrograph of green-labeled cells that have incorporated Fluo8L or neurons expressing GCaMP6f. **Middle:** Photomicrograph of red-labeled astrocytes marked with SR-101. **Right:** Overlay of the two previous photographs showing double-labeled astrocytes (arrowheads) and single-labeled neurons (arrows). The asterixis showthe positions of the pipettes used for local application of BAPTA or glutamate in the dorsal NVsnpr. **(D)** Schematic representation of the experiments depicted in **C** and **E**. **(E)** Changes in fluorescence intensity (% ΔF/F) of a neuron and an astrocyte in NVsnpr, (traces 1 and 2, respectively) and in PeriV (traces 3 and 4) in response to glutamate application (30s) in the NVsnpr.

Electrical stimulation (300 μs pulses, 40 Hz, 50-300 μA) of the dorsal part of the Vtr (**Figure 4A**) generated calcium responses in 180 cells (**Figure 4B**, 56 neurons (black empty circles) and 124 astrocytes (gray filled circles), N=14). In the NVsnpr, most neuronal and astrocytic responses were concentrated in the dorsal half with only a few cells in the ventral part (**Figure 4B, 4C and 4D**). Electrical stimulation of the Vtr also generated neuronal calcium responses in NVmt and in the area dorso-medial to it in SupV and ventral to it in PCRt. Very few neurons were responsive in the JuxtV (n=2), IntV (n=3) and NVac (n=1) **(Figure 4C** and **4E** (black bars)). Surprisingly, the distribution of activated astrocytes did not faithfully mirror that of neurons and presented some differences (**Figure 4D** and **4E** (gray bars)). Stimulation of Vtr activated more astrocytes than neurons in most areas of PeriV and dorsal NVsnpr with the highest densities found dorsally in SupV and NVsnpr. In contrast, the opposite was observed for PCRt (**Figure 4D** and **4E (**gray bars)). Three types of calcium responses were observed: repetitive transients, single transients and sustained calcium elevations as illustrated in **Figure 4F**. The bar graph in **Figure 4G** illustrates the distribution of these responses across the seven monitored nuclei for both neurons and astrocytes.

**Figure 4:**
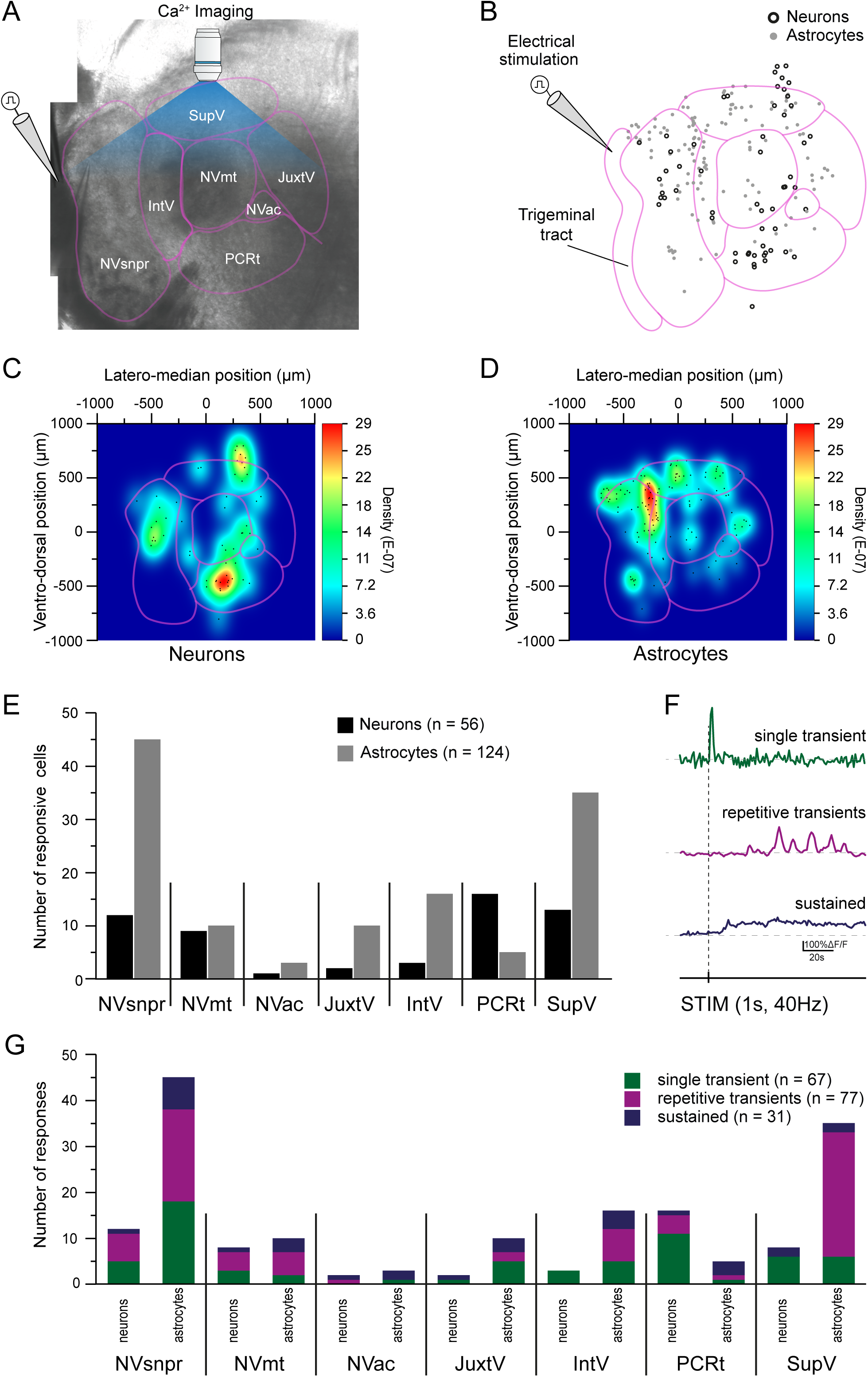
Electrical stimulation of the dorsal part of the trigeminal spinal tract generates responses in both neurons and astrocytes of the NVsnpr and PeriV. **(A)** Electrical stimulation was delivered to the trigeminal spinal tract while Ca^2+^-imaging was conducted in NVsnpr and the different areas of PeriV. **(B)** Stimulation of the Vtr elicited calcium responses in neurons (empty black circles, n=56) and astrocytes (filled gray circles, n=124) in both the NVsnpr and PeriV, as well as in the NVmt in N=14 mice. Density maps were generated for the rostral level of the brainstem (-5.34 mm to Bregma) for neurons **(C)** and astrocytes **(D)**. **(E)** The total number of responsive neurons (black bars) and astrocytes (gray bars) for each region. **(F)** Three types of calcium responses were observed in the NVsnpr and the PeriV: single transient (green), repetitive transient (magenta), and sustained (mauve). **(G)** Bar chart of the relative distribution of the three types of calcium responses in NVsnpr and PeriV.

Since NVsnpr is known to act as a relay of sensory inputs to the thalamus and perhaps other targets, and because our previous work suggests that it may form the rhythmogenic core of the CPG, we examined the distribution of cells activated when rhythmic firing is induced in its neurons by local applications of the Ca^2+^ chelator, BAPTA (as we have done before in (Morquette et al., 2015; Slaoui Hasnaoui et al., 2020; Ryczko et al., 2021; Gaudel et al., 2025)) (**Figure S2A**). BAPTA applications in the dorsal NVsnpr, elicited calcium responses in 40 neurons that were mostly concentrated in the PCRt, SupV and the dorsal part of NVsnpr (**Figure S2B** (black empty circles)**, S2C** and **S2E** (black bars), N=28). Responsive astrocytes were mostly concentrated at the junction of the NVsnpr, SupV and IntV and were sparsely distributed in the PCRt, NVmt and JuxtV (**Figure S2B** (gray filled circles)**, S2D** and **S2E** (gray bars), n=63, N=28). No neurons nor astrocytes showed sign of activation in the NVac. The same response patterns observed with the electrical stimulation of the Vtr were again identified in both neurons and astrocytes following local BAPTA application in the dorsal NVsnpr (**Figure S2F**). The bar graph in **Figure S2G** shows the distribution of these responses in the seven monitored nuclei for neurons and astrocytes.

These results confirm that electrical stimulation of the Vtr and pharmacological stimulation of NVsnpr selectively activates distinct populations of neurons and astrocytes across brainstem regions involved in sensorimotor integration. Notably, astrocytic responses were more prominent than neuronal responses in several areas, highlighting their differential spatial activation patterns and potential roles in trigeminal processing.

## Discussion

### Existence of the CMA in mice

Our study confirmed that, as in rats (Zhang and Sasamoto, 1990), there is an anterior and a posterior cortical area which stimulation drives RJMs in both awake and head-fixed animals. Optogenetic stimulation of the glutamatergic neurons of the anterior area, in the orofacial motor cortex, at similar locations to what was found by Mercer Lindsay *e*t al. (2019) in the mice, and by others in rats (Zhang and Sasamoto, 1990; Maeda et al., 2014), evoked RJMs in our animals at slightly higher frequencies to what are reported for natural mastication in mice (Kobayashi et al., 2002). As was found in rats, stimulation of the a-area, evoked masticatory movements that occurred at higher frequency and were simpler than those observed during natural mastication, as reported by Kobayashi et al. (2002), probably due to the absence of a food bolus in the mouth.

Stimulation of the p-area induces more complex, circular jaw motions with greater jaw opening in rats (Zhang and Sasamoto, 1990). This was sometimes the case in the few cases explored in this study, but further investigations are needed to compare the kinematics of the movements elicited by stimulation of the two cortical areas. Anatomically, the two regions are associated with distinct corticobulbar pathways in rats and as was found here the anterior area seems mostly related to JuxtV (Yoshida et al., 2009; Haque et al., 2010). While many similarities exist between rats and mice regarding the organization of cortical control of mastication, additional studies are needed to explore these correspondences in more detail.

### Trigeminal areas showing c-Fos activation with cortically driven RJMs or calcium-responses with electrical stimulation of the trigeminal tract The motoneuronal pools

The number of activated cells in the nuclei (NVmt and NVac) containing MNs forming the ultimate target of the masticatory command was greater in stimulated animals. No attempts were made to distinguish between MNs or interneurons, which are also part of the NVmt (Kolta, 1997; McDavid et al., 2006; Stanek et al., 2014), but in all cases, the number of c-Fos-stained cells in the NVmt was modest. Lack of sparsity of c-Fos staining, despite obvious firing activity, was also reported within the NVmt following fictive mastication (Athanassiadis et al., 2005), or in spinal MNs following fictive locomotion (Barajon et al., 1992; Dai et al., 2005). Dragunow and Faull (1989) suggested that the reason neurons in some brain regions never show c-Fos elevation despite being activated is that they probably lack the required biochemical messengers regulating Fos activation (Dragunow and Faull, 1989).

Electrical stimulation of the trigeminal tract also elicits neuronal calcium responses in the NVmt. These data are in agreement with Slaoui-Hasnaoui et al (2020) showing calcium responses in Thy1-GCaMP expressing MNs in responses to electrical stimulation of the dorsal NVsnpr which neurons receive and relay the trigeminal sensory inputs to the NVmt and surrounding nuclei (Slaoui Hasnaoui et al., 2020).

### The reticular formation

Although all subdivisions of PeriV showed significant c-Fos elevation following cortically driven RJMs, the greatest number of c-Fos-stained cells was found in JuxtV, which is still poorly investigated in mice. In the rat, it is known to receive cortical input from the CMA, to contain jaw-opening premotoneurons (Chang et al., 2009) and to mostly contribute an inhibitory input to the other subdivisions of PeriV particularly to the SupV (Bourque and Kolta, 2001) which also shows a significant number of c-Fos-stained neurons in the stimulated animals. PCRt is also strongly activated during CMA stimulation, and shares extensive interconnections with SupV, as well as with other PeriV areas. A portion of glutamatergic neurons localized in the rat PCRt receives input from SupV and NVsnpr and send their axons to NVmt and PCRtA (Kajiwara et al., 2022). In a great variety of species, such as the rat, rabbit, guinea pig and lamprey, PCRt and PCRtA contain last order interneurons to jaw muscles (Fay and Norgren, 1997; Huard et al., 1999; Kolta et al., 2000). Considering the lack of intrinsic bursting properties in this nucleus (Bourque and Kolta, 2001), the rhythmic bursts reported in PCRt during jaw-opening suggest that premotor neurons contained in this region relay the rhythm generated by other structures of the masticatory CPG to trigeminal motoneurons (Nozaki et al., 1986).

More caudally, the IRt shows relatively less c-Fos staining than in its rostral part following the cortically-induced RJMs. This area is composed of a variety of interneurons known to be involved in different orofacial behaviors, including mastication (Takatoh et al., 2021), and contains many masseter and tongue protruding premotor neurons. While tongue movements were generated during CMA stimulation, we did not observe any tongue protruding. The surprisingly smaller increase in c-Fos staining observed here may result from the absence of tongue movements due to lack of food in the mouth.

In contrast, JuxtV and IntV were rarely responsive to electrical stimulation of the trigeminal tract which instead elicit neuronal calcium responses mostly in PCRt and SupV. About 75% of SupV neurons, mainly found in its more medial part, responded to BAPTA applications in the NVsnpr as well, with a response pattern that could be associated with a rhythmic activity (sustained or repetitive) while only 30% displayed such a pattern with electrical stimulation of the trigeminal tract. This discrepancy between both modes of activation likely results from the fact that BAPTA is more prone to elicit a bursting activity in NVsnpr neurons that could be relayed as such to the connected areas. Others also reported that a proportion of SupV neurons have intrinsic rhythmogenic properties (Bourque and Kolta, 2001; Hsiao et al., 2007; Nonaka et al., 2012) and its connection with NVsnpr and NVmt raised the possibility that SupV is involved in the masticatory CPG (Hsiao et al., 2007; Nakamura et al., 2014).

### The trigeminal sensory nuclear complex (TSNC)

Mercer Lindsay et al. (2019) previously suggested that areas of the trigeminal spinal nucleus containing premotor neurons and receiving corticobulbar afferences, such as NVspo, form feedforward circuits that drive coordinated orofacial behaviors (Mercer Lindsay et al., 2019). Our current experiments did not detect significant increase in activity in NVspo in contrast to NVsnpr, which was not closely examined in the study of Mercer Lindsay et al. Surprisingly, electrical stimulation of the trigeminal sensory tract and local BAPTA applications generated only a few neuronal responses in the dorsal NVsnpr, perhaps due to the low calcium signal characteristic of the GCaMP6f (Chen et al., 2016; Gamage et al., 2023) or to the possibility that the activity of rhythmic neurons in NVsnpr does not require calcium (Brocard et al., 2006; Morquette et al., 2015) and may not be detectable with calcium imaging.

### Sensory and cortical integration

There seems to be a convergence of sensory and cortical inputs in the PCRt and at the junction of JuxtV and SupV (**Figure 5**). Those areas contain an important overlap of interneurons directly connected to sensory and cortical inputs. It is difficult to conclude how these different inputs are respectively processed. In the rat, very few rhythmic bursts were recorded in neurons from PeriV, which led us to believe that these premotor areas do not generate masticatory rhythm but may instead relay the rhythmic pattern generated elsewhere, or integrate it to other inputs before relaying it to the MNs (Bourque and Kolta, 2001). There was a sparse, but still significant, NVsnpr activation, where intrinsically rhythmic neurons are found. Given the high degree of inter-connectivity between neurons in the reticular formation and TNSC, cortical input can be rapidly distributed to all neurons in the reticular formation and TNSC. PCRt showed a strong activation in response to sensory and cortical stimulation in our experiments, which agrees with evidence, in the rat, that this area receives corticobulbar projections as well as projections from jaw muscle spindle afferents in the trigeminal mesencephalic nucleus and from SupV (Shammah-Lagnado et al., 1992; Zhang et al., 2012). Other masticatory premotor areas share extensive interconnections with PCRt (Kolta, 1997; Bourque and Kolta, 2001), and this area is often highlighted in studies on orofacial coordination. While our results do not completely elucidate the mechanisms underlying integration of these inputs, they strongly point to PCRt as an important component of the masticatory CPG for this integration.

**Figure 5:**
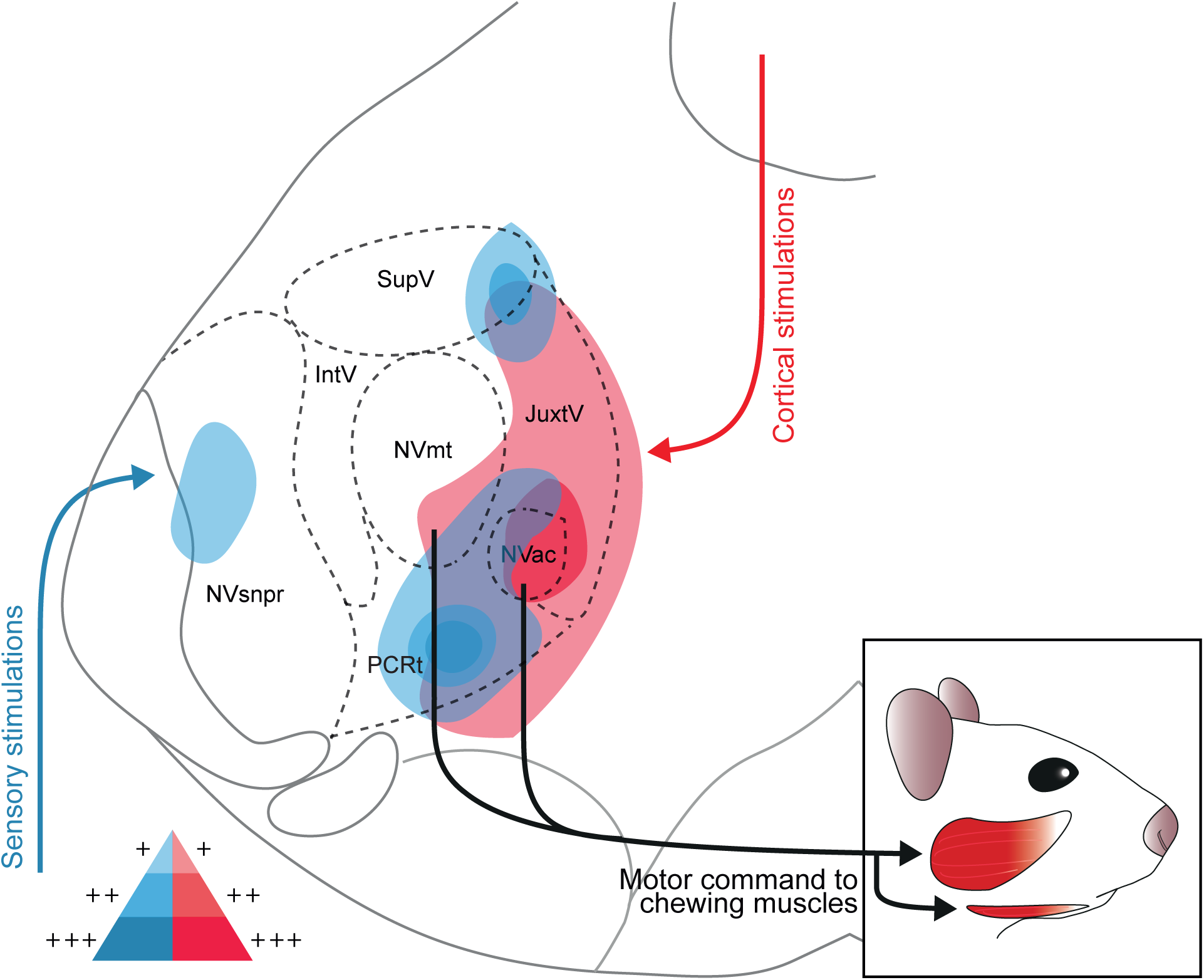
Schematic representation of the areas activated during the stimulation of cortical or sensory inputs. Stimulations of the CMA (in red) mainly engaged the JuxtV, PCRt, NVac and NVmt on both sides of the brainstem, while stimulation of the trigeminal spinal tract (in blue) engaged mainly the NVsnpr, IntV, SupV and PCRt. The areas with high saturation represent those with the highest cell density. For both types of stimulations, an overlap of activity is identified in PCRt and at the junction of SupV and JuxtV, which could indicate a convergence of cortical and sensory inputs in these areas.

## Conclusion

Mastication is more than just a preliminary step in food preparation for digestion and has been linked to important functions such as regulation of memory, vigilance, stress and anxiety (Sketchley-Kaye et al., 2011; Allen and Smith, 2012, 2015; Fukushima-Nakayama et al., 2017) emphasizing the need to better understand the circuitry underlying it.

The results presented here suggest that cortical and sensory stimulation activate distinct neuronal populations. *In vitro* experiments using brainstem slices showed that electrical stimulation of the trigeminal tract, containing the primary afferents involved in mastication, increased Ca^2+^signaling in the NVsnpr, PCRt, and SupV regions. In contrast, *in vivo* stimulation of the CMA in awake, head-fixed animals, performed here for the first time, led to a marked increase in c-Fos-labeled cells, particularly in the PeriV and PCRt regions which contain premotor neurons. These findings suggest that the PCRt may function as an integration hub for both cortical and sensory inputs. However, further studies are needed to clarify how these regions contribute to the CPG for mastication.

## Author contributions

SD, DF and OS contributed equally to data acquisition and interpretation, drafted, critically revised the manuscript and performed all statistical analyses. DV and AK contributed to conception, data interpretation, design, drafted and critically revised the manuscript. All authors have given their final approval and agreed to be accountable for all aspects of the work.

## Acknowledgements

We thank Dr. Fanny Gaudel for her contribution to the design of the figure and Yang Tian Lu for conducting two immunohistochemistry experiments presented in the paper.

## Abbreviations

CMA: Cortical masticatory area
IntV: Intertrigeminal nucleus
IRt: Intermediate reticular formation
JuxtV: Juxtatrigeminal nucleus
NVac: Trigeminal accessory motor nucleus
NVII: Facial nucleus
NVmt: Trigeminal motor nucleus
NVsnpr: Trigeminal main sensory nucleus
NVspo: Spinal trigeminal nucleus, oralis part
PCRt: Parvocellular reticular formation
PCRtA: Parvocellular reticular formation alpha
PeriV: Peritrigeminal area
SupV: Supratrigeminal nucleus
TSNC: Trigeminal sensory nuclear complex

## Supplementary Materials and Methods

### In-vivo experiments

#### Surgery and habituation

Experiments were carried out using VGluT2-Channelrhodopsin-2 (VGluT2-ChR2) transgenic mice, generated by crossing VGluT2-Cre mice (B6J.129S6(FVB)-Slc17a6^tm2(cre)Lowl^/Mwar (VGluT2-Cre), stock 028863) with ChR2-EYFP-lox mice (B6.Cg-Gt(ROSA)26Sor^tm32(CAG-COP4*H134R/EYFP)Hze^, stock 024109), at developmental stages P57-P116. Prior to the procedure, animals were anesthetized with ketamine (100 mg/kg) and xylazine (50 mg/kg) and positioned in a stereotaxic apparatus. Meloxicam (2 mg/kg) and dexamethasone (2 mg/kg) were administered subcutaneously before surgery. Bupivacaine was applied locally to the skull, which was then cleaned and drilled at the implantation site. Optic fibers were unilaterally implanted (N=4) in the a-area of the CMA (+2.5 mm AP, +2 mm L, 0.75 mm D relative to Bregma) to elicit RJMs via a 450 nm laser (Doric), secured with dental cement along with a head-bar. In another subset of mice (N=2), we bilaterally implanted in the p-area part (0 mm AP, ±3.5mm L, 2.0 mm D relative to Bregma). Anesthesia was reversed using atipamezole. Postoperative care included administration of Baytril (5 mg/kg) and buprenorphine (10 mg/kg) for pain relief, with Baytril and meloxicam continued for two subsequent days. A habituation protocol, adapted from (Guo et al. 2014), was initiated 3-7 days following surgery.

#### In-vivo stimulations and jaw movement recordings

Three to four hours before CMA optogenetic stimulation, food was removed to prevent confounding c-fos expression. Mice were head-fixed in a stabilized tube. Those implanted in the CMA received 3-second stimulations every 30 seconds for 30 minutes (40 Hz, 450 nm Doric laser, 2.5ms pulse, power: 4.54±2.20mW). Jaw movements were recorded using a 180-fps high-speed camera (Blackfly S UBS3, Teledyne Vision Solutions) controlled with MATLAB 2021A. After a 60-minute rest, the mice were anesthetized with high-dose urethane (3 g/kg) for transcardiac perfusion, starting with warm saline (0.9%) followed by cold paraformaldehyde (PFA, 4%). The brain was collected, immersed in PFA for 18 hours, then transferred to PBS.

#### DeepLabCut and frequencies analysis

CMA-induced movements were recorded using a high-speed camera and analyzed with DeepLabCut v2.3.8 (Mathis et al., 2018; Nath et al., 2019). Stimulation was identified by a LED light. We labeled 1000 frames taken from 50 videos of 11 animals and used a ResNet-50 based neural network with default parameters for 1,030,000 training iterations (He et al. 2016; Insafutdinov et al. 2016). We validated with one number of shuffles, and found the test error was: 1.67 pixel, train: 2.84 pixels (image size was 360 by 270 pixels). We then used a p-cutoff of 0.9 to condition the X,Y coordinates for future analysis. This network was then used to analyze videos from similar experimental settings. RJMs frequencies were calculated from the kinematic traces extracted from DeepLabCut and analyzed with SpinalCore software in MATLAB 2024A (MathWorks) (Mor and Lev-Tov 2007).

### In-vitro experiments

#### Brainstem and Cortical Slices Preparation

Experiments were conducted on brain slices (P8 to P18) from Vglut2-ChR2, Thy1-GCaMP6f, VGluT2-GCaMP6f, or C57BL/6 mice. Animals were anesthetized with isoflurane (Pharmaceutical Partners of Canada Inc., Richmond Hill, ON, Canada) and decapitated. Brains were quickly removed and immersed in a sucrose-based aCSF solution (in mM: 3 KCl, 1.25 KH_2_PO_4_, 4 MgSO_4_, 26 NaHCO_3_, 10 dextrose, 0.2 CaCl_2_, and 219 sucrose), cooled to 4°C, oxygenated with 95% O2 and 5% CO2, with pH 7.3-7.4 and osmolarity 300-320 mosmol/kg. Coronal slices (350 μm thick) were made using a vibratome (Leica, model VT 100S) and transferred to a room-temperature chamber filled with normal aCSF (in mM: 124 NaCl, 3 KCl, 1.25 KH_2_PO_4_, 1.3 MgSO_4_, 26 NaHCO_3_, 10 dextrose, and 1.2 CaCl_2_, pH 7.3-7.4, 290-300 mosmol/kg), oxygenated with 95% O2 and 5% CO2. Slices were allowed to rest for at least 1 hour before further experimentation.

#### Optogenetic Stimulation in Cortical Slices

CMA pyramidal neurons were optogenetically stimulated with a 440 nm laser used in the SIM light path of an FV1000 microscope (Olympus) in mice expressing the channelrhodopsin (ChR2) under the control of the Vglut2 promotor (Vglut2-Cre/ChR2-lox). Optogenetic stimulations were applied using 3 s pulses (10% laser power/8.6 µW for 440 nm laser). To assess the specificity of wavelength in activating pyramidal CMA neurons, 559 nm laser pulses (10% laser power/120 µW) were applied in control experiments.

#### Loading of Sulforhodamine-101 and calcium indicators

The brainstem slices were incubated in a holding chamber filled with normal aCSF containing 1 μM Sulforhodamine -101 (SR-101) at 35-38°C for 20 min to label astrocytes following the procedure of Kafitz et al., 2008. The slices were then transferred to a second chamber with fresh aCSF at 35-38°C for an additional 20 minutes to rinse out excess SR-101 from the tissue. Afterwards, the slices were kept at room temperature (RT) until needed. To record non-specific neuronal calcium activity, brainstem slices were incubated for 45 minutes with fluo8L-AM (20 μM, AAT Bioquest, Inc, Sunnyvale, CA, USA) and pluronic acid (20% in dimethyl sulfoxide) at RT in oxygenated aCSF, then rested for 30 minutes before data acquisition. This enabled recording activity in neurons lacking genetically encoded calcium indicators and in astrocytes.

#### Calcium imaging

PeriV is composed of SupV, JuxtV, InterV and PCRt, located dorsally, medially, laterally and ventrally to NVmt, respectively (Bourque and Kolta 2001). To map PeriV regions responding to stimulation, individual regions were monitored, and the data were superimposed onto schematic representations of the trigeminal circuit, as wide-field calcium imaging was limited by signal resolution. Calcium imaging was conducted with a confocal microscope (Olympus FluoView FV 1000) equipped with water-immersion objectives including 4X (0.28 N.A.), 10X (0.30 N.A.), and 20X (0.50 N.A.). GCaMP6f was excited with a 488 nm argon laser, with emission detected through a 500-545 nm filter. Other dyes (SR-101 and Fluo8L-AM) were excited using 559 nm and 488 nm lasers, respectively, with emissions detected through bandpass filters of 570-640 nm and 500-545 nm. Images were acquired every 0.429 second. Two-photon microscopy (BliQ) was also used, with a femtosecond laser (920 nm) and water-immersion objectives (5X or 16X), detecting SR-101, GCaMP6f and Fluo8L-AM signals through specific filters. For analysis, calcium activity was enhanced by averaging 30 images over 1 second. Topographic connectivity was assessed by recording calcium responses in NVsnpr, PeriV, and NVmt neurons as well as astrocytes following electrical stimulation of the dorsal trigeminal tract (300 μs pulse, 40 Hz, 50-300 μA) using tungsten bipolar electrodes controlled by a stimulator (Stimulus Isolator A365, World Precision Instruments) or following localized applications of 1,2-bis(o-aminophenoxy)ethane-N,N,N′,N′-tetraacetic acid tetrasodium salt (BAPTA, 5 mM, 30s) using glass micropipettes (tip diameter around 1 μm) with 2-20 psi pressure pulses (Picospritzer III, Parker Instrumentation, Fairfield NJ USA).

### Immunohistochemistry

To confirm the expression of GCaMP6f or ChR2 in Thy1 or VGluT2 neurons, brainstem and CMA brain blocks (for ChR2 mice) were sliced (40 μm thick) using a vibratome (Technical Products International Inc., Vibratome Series 1000), rinsed in PBS (5×5mins), and incubated in a blocking solution containing 0.3% Triton X-100 and 10% normal donkey serum in PBS for 1 hour. Sections were then rinsed 5×5 min in PBS and incubated overnight at 4°C in blocking solution containing primary antibodies (Rabbit anti-VGluT2, Abcam, ab229711, dilution 1:500 or Mouse anti-NeuN, Millipore, MAB377, dilution 1:600, and Chicken anti-GFP, Abcam, ab13970, dilution 1:1000). The following day, sections were rinsed 5×5 min in PBS and incubated for 90-120 min in blocking solution containing the secondary antibodies (Donkey anti-rabbit Alexa594, Jackson, 711-585-152, dilution 1:800 or Donkey anti-mouse Alexa594, Jackson, 715-585-151, dilution 1:800 and Donkey anti-chicken Alexa488, Jackson, 703-545-155, dilution 1:800). Slices were then rinsed with PBS 5×5 min before being mounted on slides using ProLong Gold antifade reagent with DAPI (invitrogen, P36935).

To confirm neuronal activation in the CMA and trigeminal circuitry after optogenetic stimulation, the same immunohistochemistry procedures were performed using primary antibodies against c-fos (Guinea-pig anti-c-fos, Synaptic Systems, 226-004, dilution 1:2000 and Mouse anti-NeuN, Millipore, MAB377, dilution 1:1000) and appropriate secondary antibodies (Donkey anti-guinea-pig Alexa Fluor 594, Jackson, 706-585-148, dilution 1:500 and Donkey anti-mouse Alexa Fluor 488, Jackson, 715-545-151, dilution 1:500).

### Images analysis

#### C-fos-positive cells analysis and heatmaps construction

Pictures were taken with a 4X magnification and cells were counted automatically by ImageJ Fiji for each nucleus according to the schematic illustrations in Figure 2. Nuclei identification was conducted by the same person on all brain slices with pictures revealing only NeuN antibody to avoid the bias brought by the presence of c-fos-stained cells. Using images with NeuN staining to align nuclei at the same coordinates, images of c-fos staining were taken from each mouse. Coordinates of the cells were calculated by FIJI ImageJ and were fed in OriginPro 2022b software to automatically generate cell density maps.

#### Calcium imaging analysis and mapping procedures

Fluorescence intensity changes were calculated as %ΔF/F0 = [(F − F0)/F0] × 100, with ROIs drawn around cell bodies. The dFoverFmovie plugin in ImageJ helped detect weaker responses obscured by background fluorescence. Cells with fluorescence increases ≥50% were included. Plateau responses lacked distinguishable oscillations, while repetitive transient responses showed clear oscillations. Their frequency was analyzed after smoothing the signals using the Savitzky-Golay method in OriginPro 2022b.

Coordinates (x; y) of responsive cells to the electrical stimulation of the dorsal trigeminal tract (Vtr) were mapped and graphically represented in normalized schemas of NVsnpr, NVmt, and PeriV (Condamine et al. 2018). To standardize all data, coordinates were scaled to values between 0 and 1 by dividing by the image’s total length (x-axis) or width (y-axis).

High-resolution images (16X, 20X) were resized to align with low-magnification references (5X, 4X) in Inkscape 1.1.2 software, ensuring coordinates could be adjusted. By aligning the top-left corner of the resulting image to the origin, the cells recorded coordinates at 16X or 20X could be read in the 5X and 4X references, denoted as **(x_5X_; y_5X_ and x_4X_; y_4X_**, respectively). Using FIJI ImageJ, reference nuclei were outlined with a Bounding Rectangle to measure their dimensions **(w, h)** and positions **(x_BR_; y_BR_)**. Cell coordinates within the rectangle were calculated as **(x_R_; y_R_) = (x_5X_ - x_BR_; y_5X_ - y_BR_)**, where **(x_R_; y_R_)** are the coordinates of the cell within the rectangle defined by the ROI; **(x_5X_; y_5X_)** are the coordinates of the cell in the 5X reference; and **(x_BR_; y_BR_)** are the coordinates of the Bounding Rectangle reference. This yielded **(x_R_; y_R_)**, the coordinates representing the cell’s position along the x and y axes in the schematic representation of the artificial nucleus.

The PeriV cells coordinates were obtained using NVmt as the reference nucleus, and to apply the protocol just described, images were arbitrarily oriented so that NVsnpr was on the left side of the image, and the midline was on the right side of the image. To generate cell density maps, the x- and y-coordinates of the cells were used in OriginPro 2022b software.

#### Statistics and graphics

Data are presented as mean ± SEM. “N” denotes animals and “n” the cells analyzed Statistical tests were performed using IBM SPSS Statistics and GraphPad Prism 10.5.0 included two-tailed Wilcoxon matched-pairs signed rank test for frequencies analysis and Mann-Whitney U tests due to non-normality distributed data for c-Fos data with Holm-Šídák correction for multiple comparisons. Significance was set at P < 0.05.

**Figure S1:**
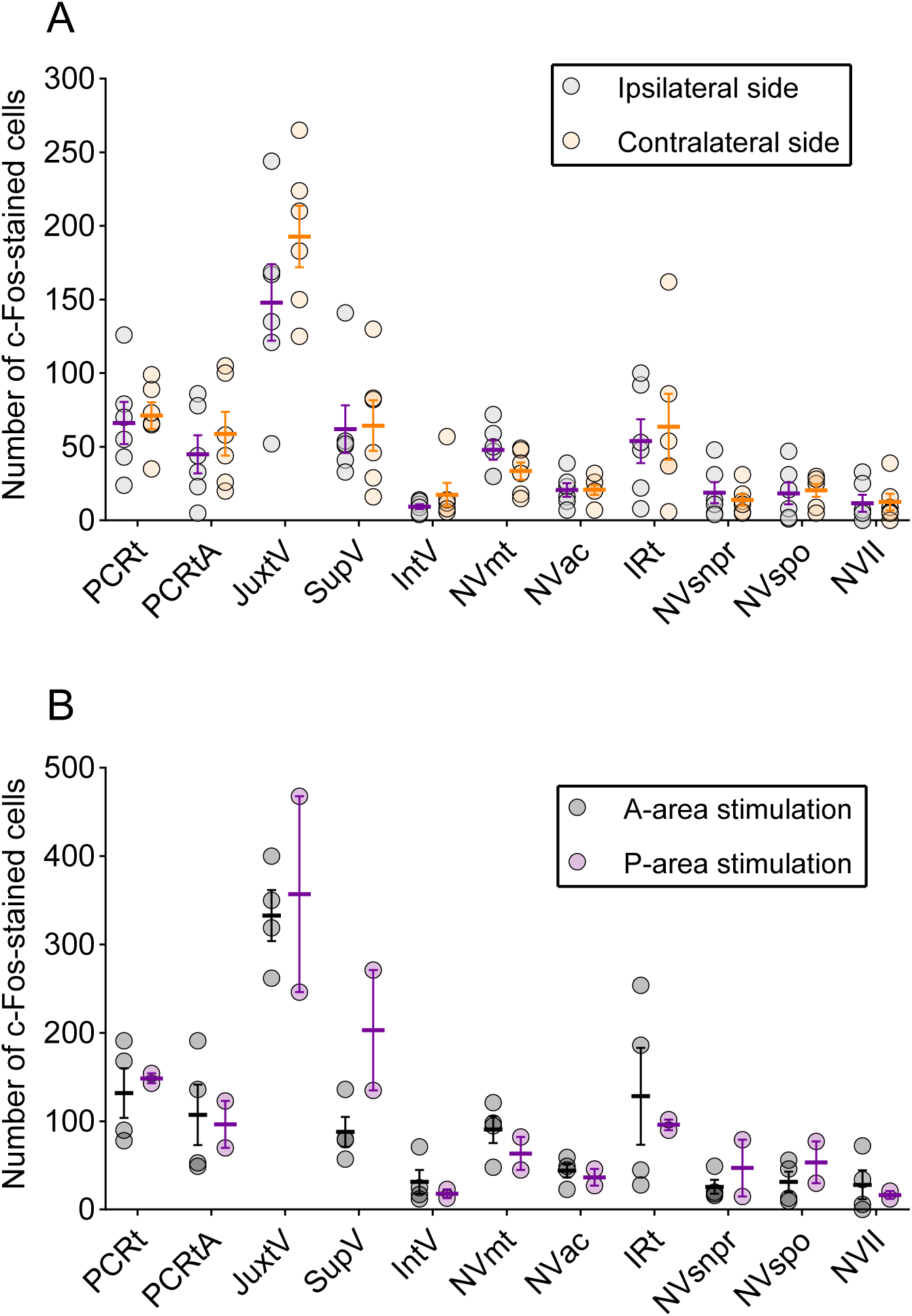
The side and site of stimulation have little effect on the number of c-Fos-stained cells. **(A)** Quantification of c-Fos-stained cells relative to the lateralization of the stimulation site showed no significant difference between the ipsilateral and contralateral sides. **(B)** Stimulations in the a-area and p-area regions did not lead to any significant difference in the number of c-Fos-stained cells. All observed *P*-values are above 0.5.

**Figure S2:**
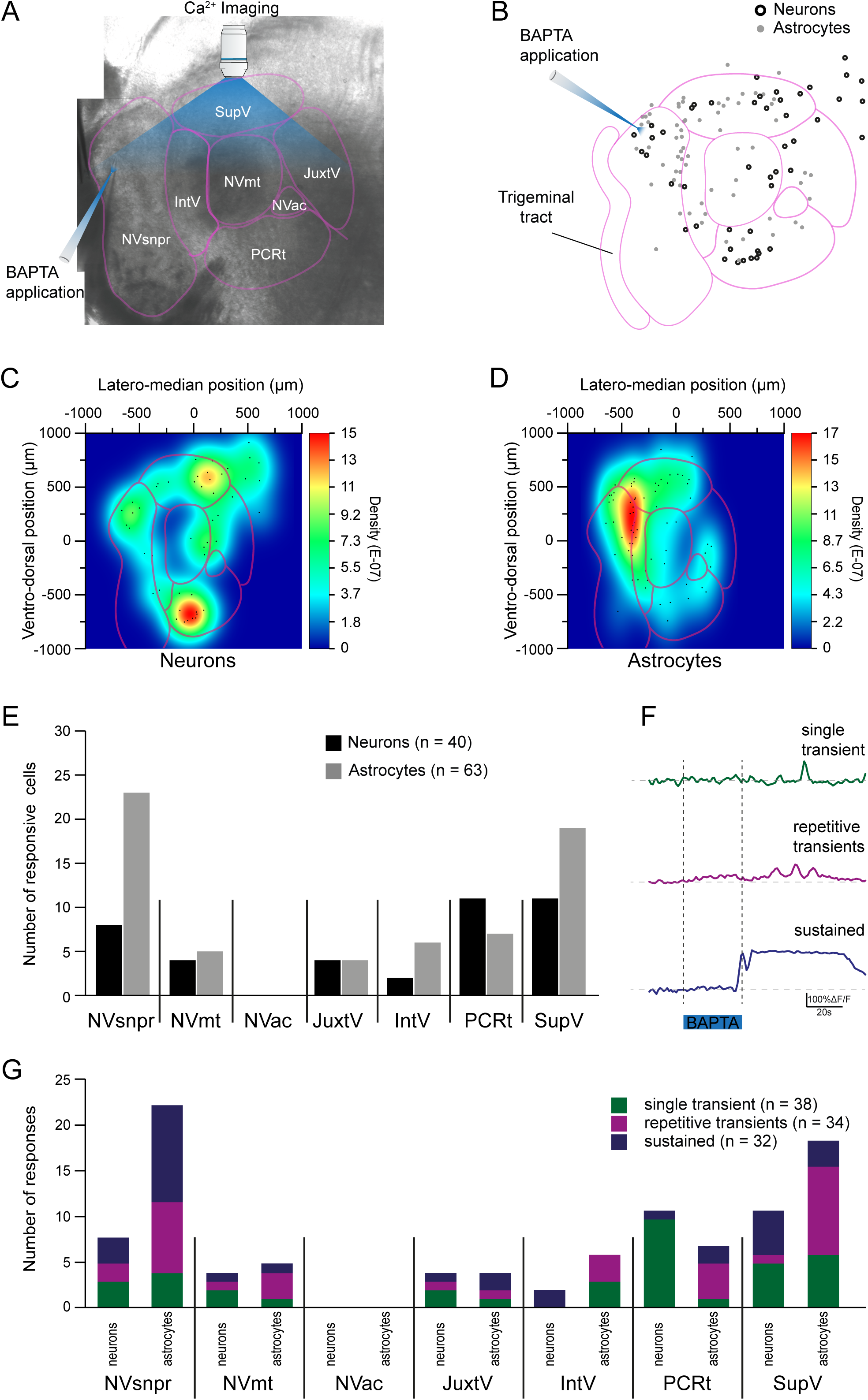
BAPTA applications in the dorsal part of NVsnpr generate rhythmic responses in both neurons and astrocytes of the NVsnpr and PeriV. **(A)** BAPTA applications were delivered for 25.4 ± 1.4 seconds in the NVsnpr while Ca^2+^ imaging was conducted in NVsnpr and the different areas of PeriV. **(B)** BAPTA applications elicited calcium responses in neurons (empty black circles, n=40) and astrocytes (filled gray circles, n=63) in both the NVsnpr and PeriV, as well as in the NVmt in N=28 mice. Density maps were generated for the rostral level of the brainstem (-5.34 mm to Bregma) for neurons **(C)** and astrocytes **(D)**. **(E)** The total number of responsive neurons (black bars) and astrocytes (gray bars) for each region. **(F)** Three types of calcium responses were observed in the NVsnpr and the PeriV: single transient (green), repetitive transient (magenta) and sustained (mauve). **(G)** Bar chart of the relative distribution of the three types of calcium responses in NVsnpr and PeriV.

